# Open Digital Bioassays Enabled by Aqueous Two-Phase Microreactor Arrays

**DOI:** 10.64898/2026.07.12.738022

**Authors:** Yoshihiro Minagawa, Kai Matsumoto, Shoki Nakata, Hideaki Isago, Masaomi Nangaku, Makoto Kurano, Hiroyuki Noji

## Abstract

Digital bioassays enable precise single-molecule quantification but are difficult to adapt to point-of-care testing (POCT) because conventional protocols include off-chip complex processes for sample treatment and sealing, requiring hardware and workflow complexity. We present OASSIS (Open Aqueous two-phase Separation System for Integrated Single-molecule digital bioassay platform), an oil-free and open ATPS platform that localizes both targets and signals in femtoliter-scale dextran (DEX) droplets beneath a polyethylene glycol (PEG) phase. We integrated a CRISPR-Cas13a assay system with a novel, branched fluorescent reporter conjugated to a dextran-binding domain (DBD), which ensures signal retention within the DEX droplets after cleavage. Fluorescence recovery after photobleaching experiments confirmed this robust signal confinement. OASSIS not only performs amplification-free digital RNA detection but also enables serial sample introductions through its open-format architecture that progressively improve sensitivity: the limit of detection (LOD) improved from 1.08 fM (first introduction) to 0.34 fM (third introduction). Furthermore, OASSIS demonstrated specific detection and ∼10-fold enrichment from a complex, denaturant-treated nasopharyngeal swab matrix. Together, these results demonstrate that the open-format architecture of OASSIS provides a practical route toward sensitive, low-complexity POCT and clinical diagnostic applications.

## Introduction

Digital bioassay is a single-molecule detection format designed to quantify individual target molecules with high precision^1–6^. In this method, the assay mixture is divided into a large number of micrometer-scale reactors, leading to a stochastic distribution of target molecules among them. Each reactor produces an optical signal, typically fluorescence, from enzymatic reactions catalyzed by the target molecule or a bound marker enzyme. After binarization of the signal (i.e., classifying reactors as positive or negative based on fluorescence intensity, 0 or 1), the absolute target count is estimated from the fraction of positive reactors according to a Poisson occupancy model. In conventional digital bioassays, water-in-oil emulsion systems are employed for micro-compartmentalization. Such approaches, exemplified by digital PCR, LAMP, and ELISA, have established digital bioassay as a powerful analytical technology for precise quantification and rapid diagnostics^7–11^. Although various types of micro-compartmentalization^10, 12^ and compartmentalization-free methods^13–17^ were reported, the translation of digital bioassays into point-of-care testing (POCT) remains challenging.

A key limitation for POCT is its reliance on hardware-based sample enrichment ^18–20^. Enrichment of rare targets from dilute samples typically requires centrifugation or magnetic concentration after capturing target molecules on magnetic beads, increasing operational complexity and limiting the practicality of these methods outside centralized laboratories ^9, 10, 21–23^. Therefore, hardware-free enrichment is essential for the broad deployment of POCT technologies. To address this, we exploited the properties of an aqueous two-phase system (ATPS) composed of dextran (DEX) and polyethylene glycol (PEG), which spontaneously enriches various biomolecules in DEX-rich droplets ^24–27^. ATPS of DEX/PEG system and relevant systems were used for various bioanalytical methods ^28–31^. We developed a femtoliter reactor array device (FRAD) that forms micron-sized, regularly shaped DEX droplets beneath the PEG solution ^32^. These DEX droplets enabled highly sensitive digital assays, actively enriching nucleic acids and proteins tagged with a dextran-binding domain (DBD) derived from dextransucrase from *Leuconostoc mesenteroides* ^33^.

Another challenge lies in the closed nature of microreactors. The reactor sealing process itself introduces procedural complexity and restricts the effective sample processing volume. Oil sealing is conventionally used for micro-compartmentalization in both flow-through water-in-oil emulsion systems and droplet array systems including FRAD. Recently, alternative oil-free methods have been reported, including tape/film-based sealing and air sealing; however, these methods still require discrete sealing steps, adding procedural complexity^34–36^. In addition, once sealed, these micro-compartmentalization methods restrict the assay to the finite sample volume loaded into the reactors, which can be limited to nanoliter-scale effective volumes in reactor-array formats, leaving much of the collected specimen unused as dead volume.

This study aims to develop a new digital bioassay strategy—Open Aqueous two-phase Separation System for Integrated Single-molecule digital bioassay platform (OASSIS)—that enables the enrichment of target molecules and the signal localization without a sealing process. This novel approach alleviates the limitations of restricted input volume by allowing serial sample introduction. Building upon the DEX droplet-array system, we integrated the Cas13-based RNA detection assay to demonstrate the concept. For the spatial localization of the signal fluorescence generated by Cas13 enzyme, we anchored a fluorescent reporter to the DEX phase via DBD tag. Using this configuration, we performed stepwise target additions to the same array and observed monotonic sensitivity gains with cumulative sample volume. OASSIS eliminates the need for reactor sealing, maintaining binary quantification while alleviating constraints of traditional closed systems.

## Results

To establish a model digital bioassay with the Open Aqueous two-phase Separation System for Integrated single-molecule digital assay platform (OASSIS), we employed a CRISPR-Cas13-based digital RNA detection system ^32, 37, 38^. The working principle of the RNA digital detection on OASSIS is illustrated with that for the conventional method (Fig. 1a, b). Upon binding to a target RNA via its guide crRNA, Cas13 protein activates a non-specific trans-RNase activity and cleaves self-quenched reporter RNA probes, yielding fluorescence signal (Fig. 1a) ^39, 40^. Typical length of RNA probe is about 6-mer, which is insufficient for partition in the DEX-rich phase that requires 100 nt or more ^41^. Therefore, in our previous work with DEX droplet reactors on femtoliter array device (FRAD), the reactors had to be sealed with oil to retain fluorescence probe molecules in the droplets (Fig. 1c). For the open-format digital bioassay platform, Cas13 enzyme protein and RNA probe are designed to be entrapped in DEX droplets (Fig. 1b); each of Cas13 protein (*Lwa*Cas13a) and RNA probe was tagged with a tandem repeat of dextran-binding domain protein (DBD×2) for spontaneous enrichment (Fig. 1b and 1d). In addition, the RNA probe was designed to have a branched structure so as to retain the conjugation with fluorophore with DBD×2 after cleavage from the quencher moiety. The reporter design enables the enrichment of RNA probe molecules in DEX droplets, allowing oil-free, open-format detection.

**Figure 1.**
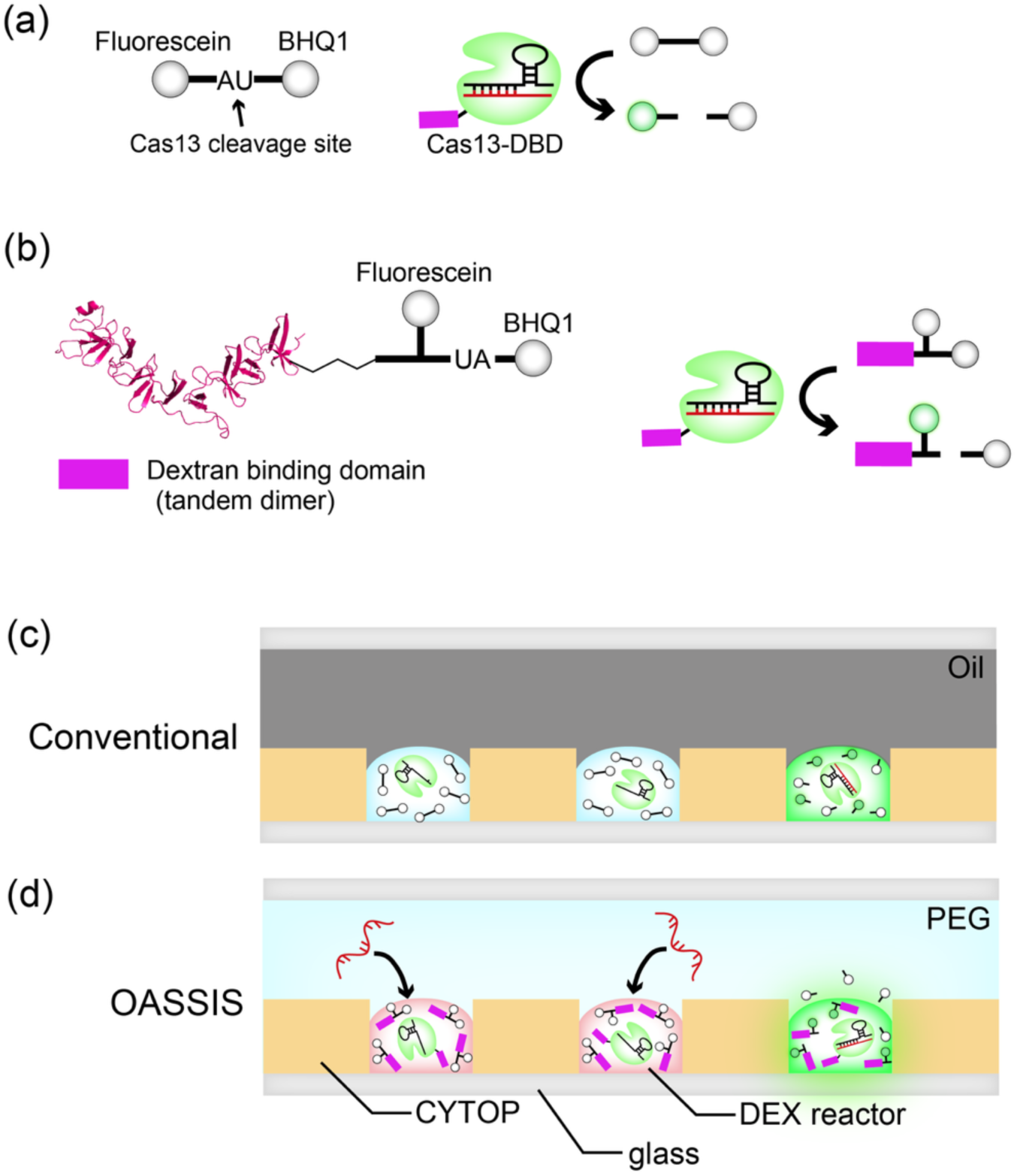
Concept and key components of OASSIS. **(a)** Schematics of the reporter probe and the reaction scheme of Cas13 assay. Cas13 is tagged with a tandem dextran-binding domain, DBD×2 (magenta) for spontaneous enrichment in DEX reactor. Upon target RNA recognition, Cas13a cleaves the probe at the indicated site, generating a fluorescent signal. **(b)** Schematics of the branched reporter probe and the reaction scheme. The branched nucleotide strand is conjugated to DBD×2 (magenta), a fluorophore (Fluorescein, F), and a quencher (BHQ1, Q). The branched structure ensures that the fluorophore remains anchored to the DBD×2 and is retained within the DEX reactor. **(c)** Schematics of the conventional oil-sealing digital bioassay with DEX reactor system, used as a control assay. **(d)** Schematics of OASSIS. DEX reactors form an open aqueous interface with the overlaying PEG (polyethylene glycol) phase, enriching target RNA molecules from the PEG phase into the DEX reactors, while DEX reactors retain the DBD-conjugated reporter molecules cleaved by Cas13, generating fluorescence signal.

To test the possible cross talk of the RNA probe among DEX droplets, we conducted a fluorescence recovery after photobleaching (FRAP) experiment, using a model RNA-DBD conjugate; the branched reporter nucleotide strand conjugated with DBD×2 (Fig. 2a). After the fluorescent probe was enriched from the PEG-rich phase in the DEX droplet array, target reactors were photobleached using focused laser spots. The subsequent fluorescence recovery was monitored over time (Fig. 2b). Images were acquired before and after photobleaching; 0, 10, 20, and 30 min (Fig. 2c). Fluorescence recovery was not observed throughout the 30-minute observation period (Fig. 2d). The slow decrease in fluorescence of non-bleached reactors was due to photobleaching. These observations show that the leakage of DBD-conjugated probe among reactors is practically negligible, confirming robust enrichment of the reporter probe in DEX droplet reactors in the open format. Similarly, FRAP analysis confirmed that Cas13-DBD was also stably retained in DEX droplets under open-format conditions (Supplementary Fig.1).

**Figure 2.**
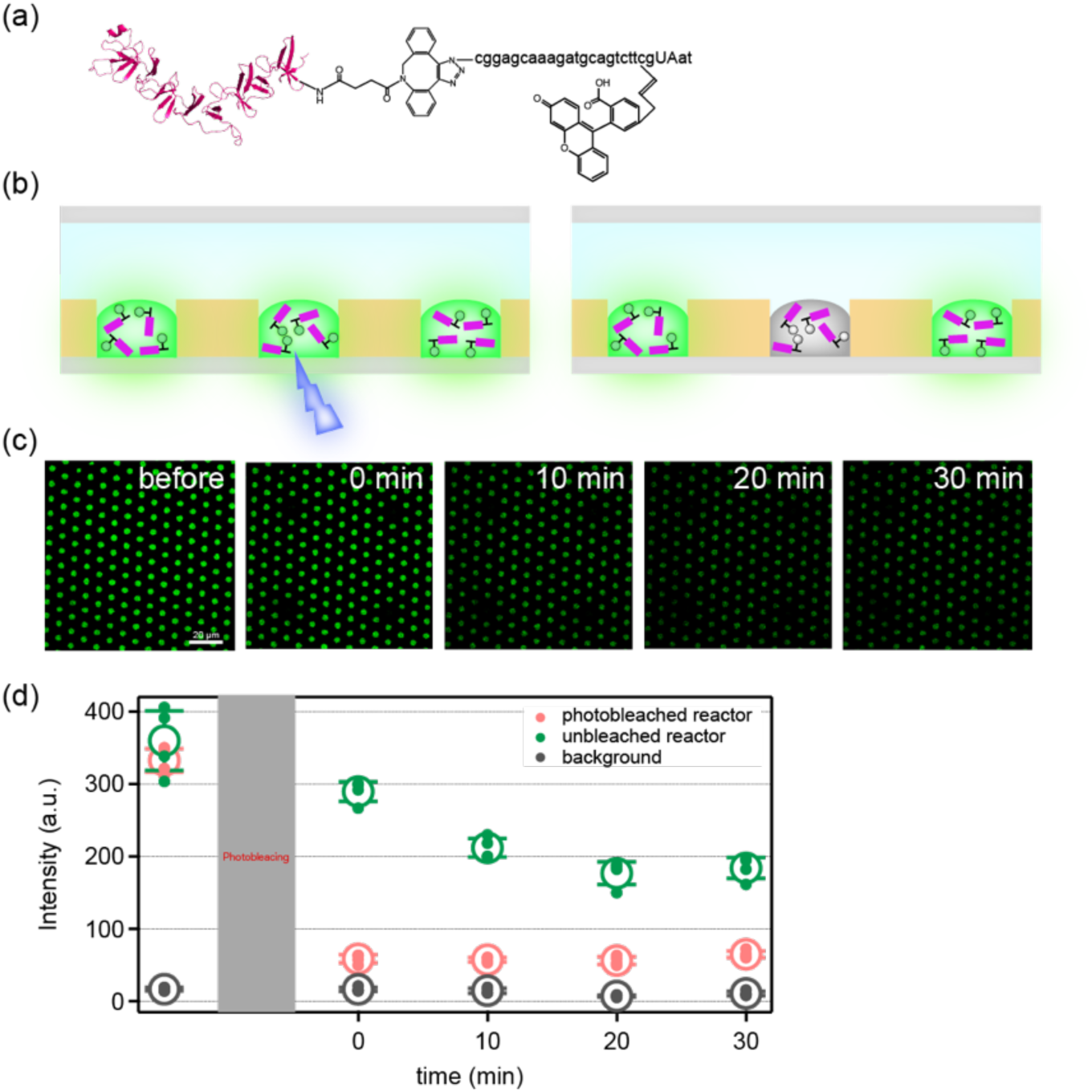
FRAP analysis of DBD-conjugated reporter probe. **(a)** Structure of the model DBD-conjugated fluorescent probe, mimicking DBD-conjugated branched reporter probe after Cas13 cleavage. **(b)** Schematic of the Fluorescence Recovery After Photobleaching (FRAP). **(c)** Time-lapse fluorescence images of FRAP analysis. Scale bar, 20 μm. **(d)** Time-course of fluorescence intensity of photobleached reactors (red circles), neighboring unbleached reactors (green circles), and the background (gray circles). Circles and error bars represent the mean ± s.d. (*n* = 4)

Having validated the robust retention of the DBD-conjugated probe, we next evaluated the performance of the complete seal-free OASSIS for digital RNA detection. For comparison, a conventional oil-sealed digital assay using the same Cas13/crRNA system was performed in parallel (Fig. 3a). In OASSIS, the assay was initiated by forming DEX reactors and enriching them with Cas13-DBD, crRNA, and the DBD probe. A PEG solution containing target RNA was subsequently introduced into the flow channel, and the reaction was monitored by time-lapse fluorescence imaging (Fig. 3c).

**Figure 3.**
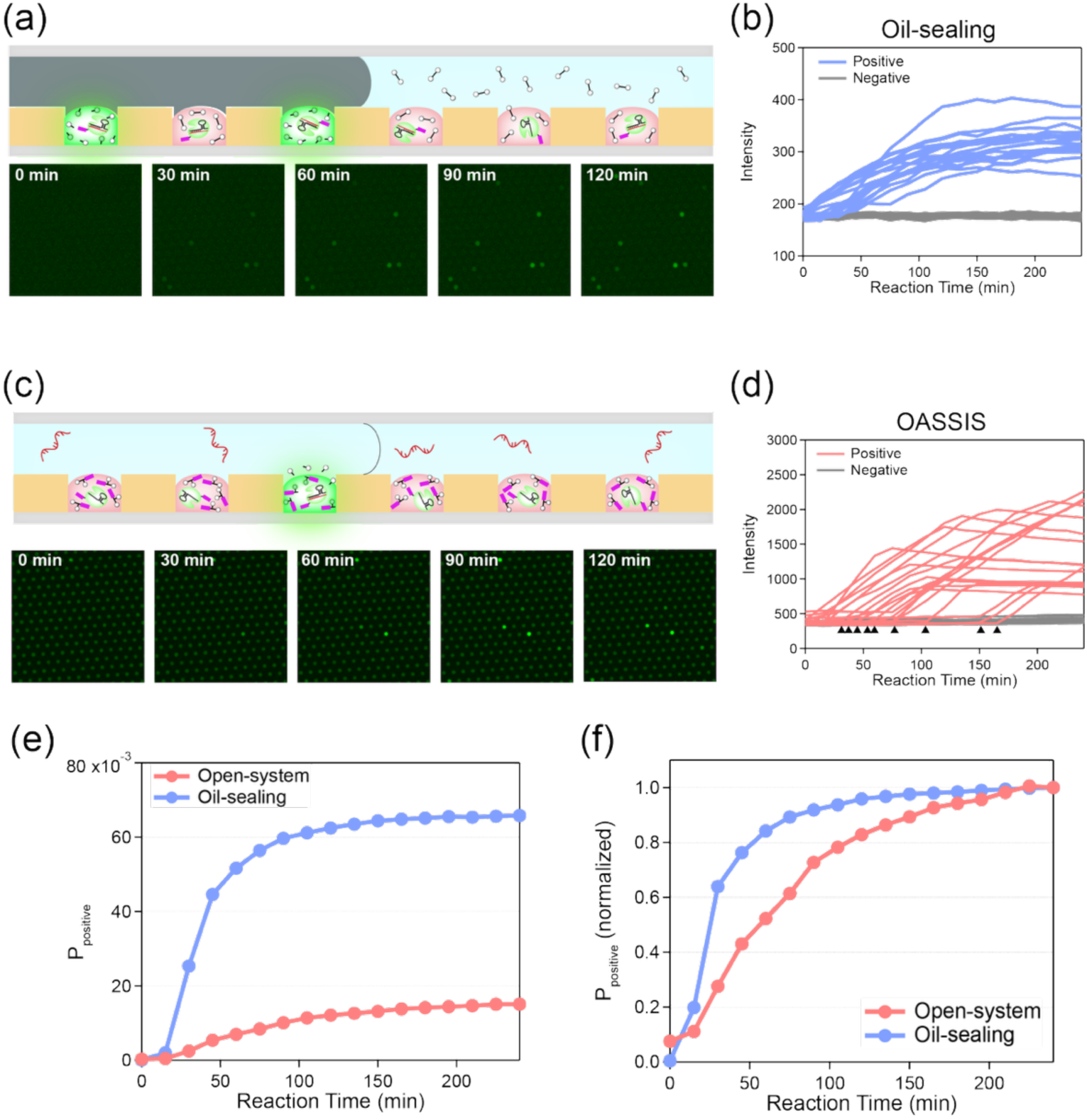
Digital RNA detection with the oil-sealed system or with OASSIS. **(a)** Schematic and time-lapse fluorescence images of the Cas13 digital bioassay in the conventional oil-sealed DEX droplet system. **(b)** Time-courses of positive or negative reactors found in oil-sealed DEX droplet system. **(c)** Schematic and time-lapse fluorescence images of the assay on OASSIS. **(d)** Time-courses of positive or negative reactors found in the assay on OASSIS. **(e)** Temporal evolution of the probability of positive reactors (*P*_positive_) in OASSIS (red) and the oil-sealed system (blue). The final plateau level of OASSIS was approx. 1/4 of the closed system. (f) Normalized *P*_positive_ time-courses. The half-times (T_1/2_) for OASSIS (red) and the oil-sealed system (blue) were approx. 56 min and 25 min, respectively.

Representative fluorescence time-courses of individual reactors classified as positive or negative are shown in Fig. 3b,d. In both the oil-sealed system and OASSIS, positive reactors exhibited a lag phase followed by a rapid increase in fluorescence intensity, whereas negative reactors remained at baseline levels throughout the observation period. Notably, in OASSIS, some positive reactors displayed substantially longer lag times before the onset of fluorescence increase (Fig. 3d). At first glance, this behavior could be interpreted as a consequence of delayed transport of target RNA molecules from the surrounding PEG phase into the DEX reactors. However, similar delayed activation events were also observed in the conventional oil-sealed system (Supplementary Fig. 2b), where neither external target supply nor inter-reactor transport can occur. This observation is consistent with the possibility that the branched reporter architecture influences signal-generation kinetics.

We next analyzed the temporal evolution of the positive-reactor probability (*P*_positive_), defined as the fraction of reactors exceeding the fluorescence threshold (Fig. 3e). A clear difference was observed between the two systems. In the oil-sealed assay, *P*_positive_ increased rapidly and reached a plateau within approximately 100 min. In contrast, OASSIS exhibited a more gradual increase in *P*_positive_ over time. Consequently, the plateau value of *P*_positive_ in OASSIS was substantially lower than that of the oil-sealed system, reaching approximately one-quarter of the final levsel observed in the conventional assay. A similar reduction in the final *P*_positive_ was also observed in the oil-sealed assay using the DBD-conjugated branched reporter (Supplementary Fig. 2c), suggesting that the reduced plateau was associated with the branched reporter design rather than the open-format configuration itself.

To compare the reaction kinetics independently of the final positive-reactor fraction, *P*_positive_ values were normalized to their respective plateau values (Fig. 3f). After normalization, the temporal profiles of the two systems became considerably more similar. The half-times (*T*_1/2_) of OASSIS and the oil-sealed system were approximately 56 min and 25 min, respectively. These results suggest that the principal difference between the two systems lies in the final fraction of reactors that become positive, rather than in the fluorescence activation kinetics of individual positive reactors.

We then tested serial sample injection in OASSIS, taking advantage of its open-format architecture (Fig. 4a). After each round of sample injection, the fluorescence images were taken (Fig. 4b). The number of the positive reactors significantly increased upon serial sample introduction (Fig. 4b). For quantitative analysis, the assay was conducted at different target RNA concentrations (Fig. 4c). *P*_positive_ increased with the target RNA concentration, showing clear linear response. Notably, owing to the on-chip enrichment capability of OASSIS, *P*_positive_ after 1st sample injection (red circles in Fig. 4c) already exceeded the theoretical value (gray line in Fig. 4c) estimated without consideration of enrichment effect. The limit of detection (LOD) after the first sample injection was determined to be 1.08 fM. Upon additional sample injections (the second and third injections in Fig. 4c), *P*_positive_ significantly increased. Accordingly, the LOD was also improved, decreasing to 0.53 fM after the second injection and 0.34 fM after the third injection, thereby reaching the sub-femtomolar range. Thus, it was confirmed that the open nature of OASSIS allows serial sample injection to enhance the effective sample volume and detection efficiency.

**Figure 4.**
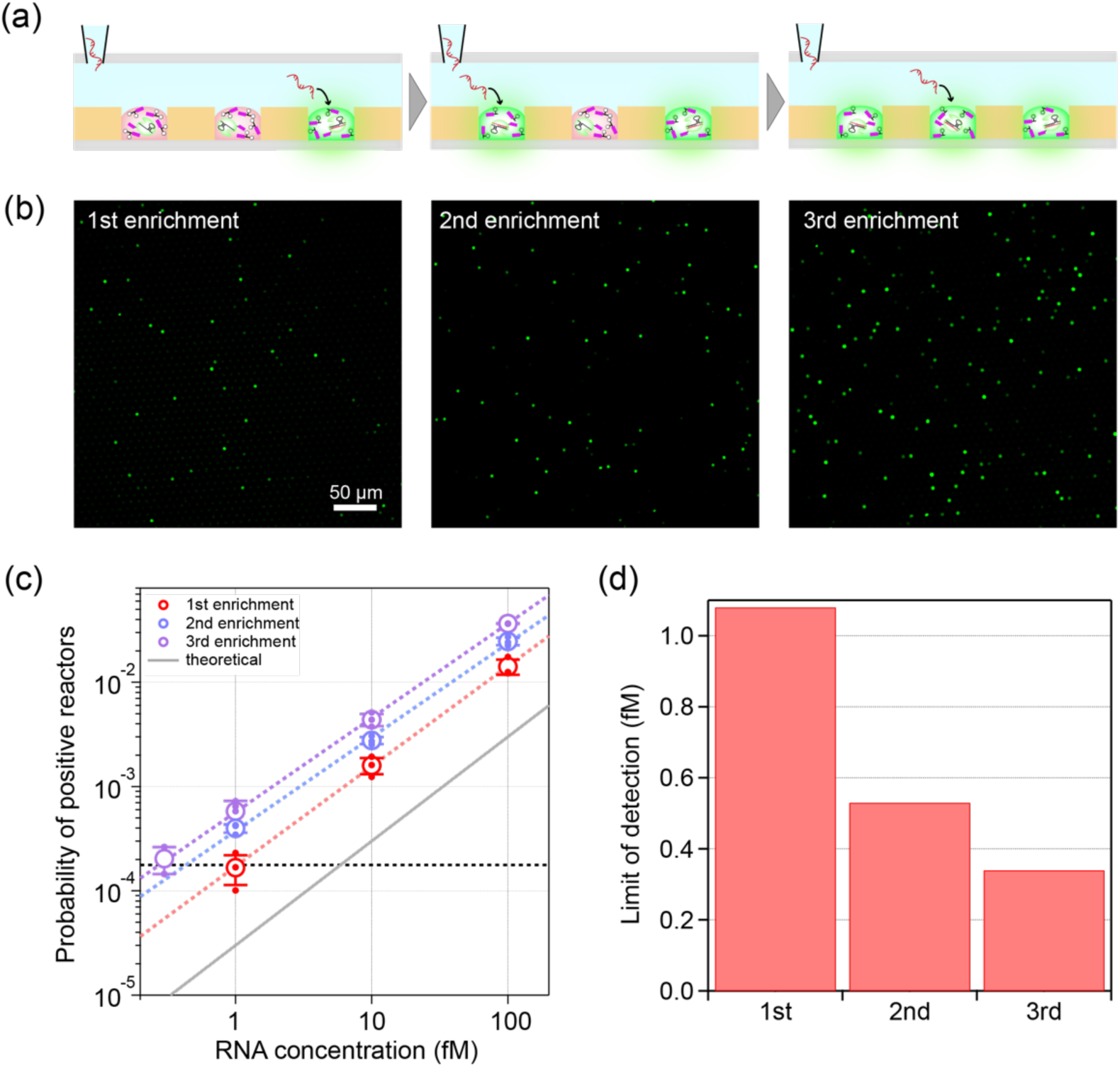
Serial sample enrichment for enhanced detection sensitivity. **(a)** Schematic illustration of serial sample introduction. Target RNA molecules are sequentially introduced from the PEG phase. **(b)** Fluorescence images of the DEX reactor array after the 1st, 2nd, and 3rd rounds of sample introduction (enrichment). Scale bar, 50 µm. **(c)** *P*_positive_ against target RNA concentration after the 1st (red circles), 2nd (blue circles), 3rd (purple circles) sample injections, and the theoretical values estimated from RNA concentration in PEG phase without considering enrichment by DEX (gray solid line). Black dashed line indicates the limit of detection (LOD). Circles and error bars represent the mean ± s.d. (n=3). **(d)** LOD after the first, second, third rounds of sample injection.

Finally, we investigated the practical utility of digital RNA detection with OASSIS for clinical applications, by conducting a spike-in experiment using a nasopharyngeal swab sample from a healthy donor. A primary challenge for the assay with clinical specimens is the degradation of RNA molecules by endogenous RNase in the sample. In standard assays for retrovirus RNA detection, denaturation reagents, such as guanidinium thiocyanate (GITC) and 2-mercaptoethanol (2-ME), are often used for the virus lysis and the inhibition of endogenous RNase activity. However, because the denaturation reagents also interfere with downstream enzymatic assays ^42^, subsequent purification process is required. We found that the standard denaturing condition used here (1 M GITC and 1% 2-ME) almost completely inhibited Cas13 activity. In contrast, Cas13 retained its enzymatic activity when the denaturant-treated sample was diluted 200-fold, corresponding to final concentrations of 5 mM GITC and 0.005% 2-ME (Supplementary Fig. 3).

Based on this finding, we prepared a simulated clinical sample by spiking the target RNA into the denaturant-treated nasopharyngeal swab matrix, which was then diluted 200-fold with the PEG solution for the assay (Fig. 5a). The target RNA was spiked in at a final concentration of 300 fM after dilution. At this concentration, the theoretical fraction of positive reactors without the consideration of enrichment effect by DEX droplet is 0.01. The assay revealed a clear distinction between the spiked sample and a negative control: the positive fraction for the spiked sample was 0.10, whereas the negative control yielded only 3.8×10⁻⁵ (Fig. 5b, 5c). This empirically measured positive fraction of 0.10 is 10-fold higher than the theoretical prediction. This result demonstrates that our platform efficiently concentrates the target molecule approximately 10-fold and enables specific detection from a denaturant-treated clinical sample matrix.

**Figure 5.**
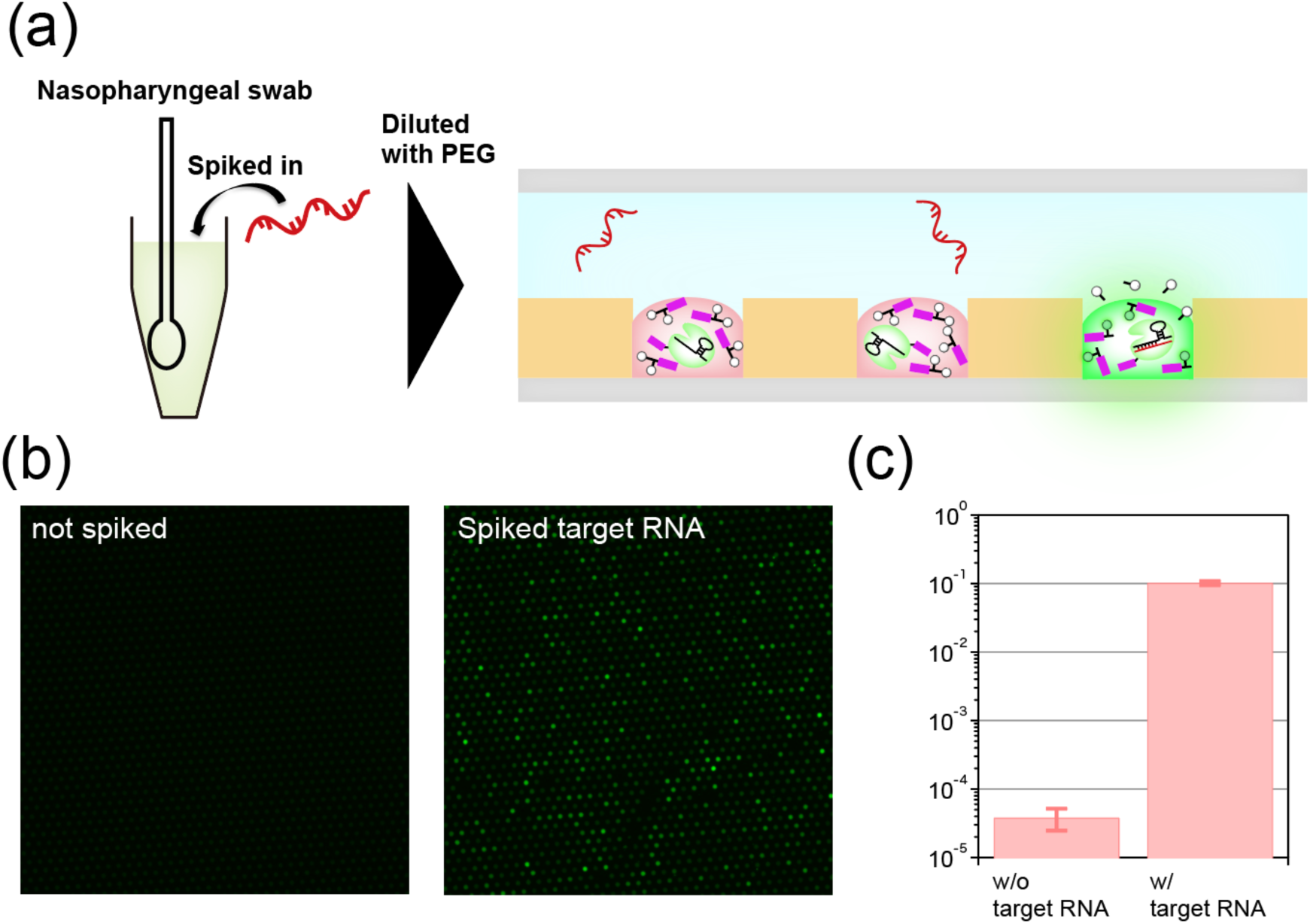
Digital RNA detection of a complex clinical matrix. **(a)** Schematic of the spike-in experiment. Target RNA was spiked into a denaturant-treated nasopharyngeal swab sample from a healthy donor. The sample was then diluted 200-fold with the PEG solution and introduced into OASSIS. **(b)** Representative fluorescence images of the reactor array for the negative control (not spiked, left) and the sample spiked with target RNA (right). The final concentration of spiked RNA after dilution was 300 fM. **(c)** Quantification of the probability of positive reactors (*P*_positive_) for the sample without target RNA and with target RNA. Error bar represents s.d. (*n* = 3).

## Discussion

In this study, we developed OASSIS (Open Aqueous two-phase Separation System for Integrated Single-molecule digital bioassay platform) by integrating a newly designed DBD-conjugated reporter probe with a previously established dextran droplet array system. This approach enabled amplification-free digital RNA detection based on CRISPR-Cas13 in a completely oil-free format. OASSIS achieved a limit of detection of 1.08 fM from a single sample introduction, which was further improved to 0.34 fM through three rounds of serial sample introduction. In addition, we demonstrated specific detection of target RNA from a denaturant-treated nasopharyngeal swab matrix, confirming the compatibility of the platform with clinically relevant samples. These results establish OASSIS as a new digital bioassay architecture that combines molecular enrichment, digital quantification, and open-format operation in a single platform.

The most distinctive feature of OASSIS is its ability to perform serial enrichment through repeated sample introduction. In conventional digital bioassays, the probability of target detection is fundamentally determined by the concentration of target molecules within a fixed sample volume. In contrast, OASSIS allows target molecules to be continuously accumulated into the same reactor array over multiple introduction steps. As a result, the probability of detection becomes increasingly dependent on the total number of target molecules processed rather than solely on their initial concentration. This concept expands the conventional framework of digital bioassays by shifting the determinant of detection sensitivity from target concentration to the total number of target molecules processed. As such, OASSIS may be particularly advantageous for applications involving rare targets distributed in large sample volumes, including environmental monitoring, food safety testing, and infectious disease diagnostics.

The open architecture of OASSIS also addresses one of the major practical limitations of conventional digital assays: the requirement for reactor sealing. In most existing digital platforms, compartmentalization requires oil sealing or other physical isolation procedures, which increase workflow complexity and restrict the effective sample volume that can be processed. By localizing both target molecules and fluorescence signals within DEX droplets, OASSIS maintains digital quantification without a sealing step. Furthermore, the platform successfully detected target RNA in a denaturant-treated nasopharyngeal swab matrix after simple dilution, demonstrating compatibility with a clinically relevant sample environment. Although additional developments will be required for practical point-of-care implementation, including direct handling of crude samples and integration of simplified sample preparation procedures, the present results demonstrate a promising route toward low-complexity digital diagnostics.

Despite these advantages, one important limitation of the current implementation of OASSIS became apparent. OASSIS exhibited a substantially lower plateau value of *P*_positive_ than the conventional oil-sealed assay. The time-course analysis provides an important clue to understanding this behavior. Many positive reactors exhibited prolonged lag phases before the onset of fluorescence increase (Fig. 3d). This observation suggests that signal generation is substantially delayed in at least a subset of reactors. Consequently, some reactors containing target RNA may not have reached the fluorescence threshold within the observation period and therefore remained classified as negative, leading to an underestimation of the final positive-reactor fraction.

The reduced plateau level is attributed to the branched DBD-conjugated reporter used in the present study. Supporting this interpretation, control experiments demonstrated that the branched reporter markedly reduced the final positive-reactor fraction even in the conventional oil-sealed DEX droplet assay (Supplementary Fig. 2). This result suggests that the reporter design, rather than the open nature of the system, is the major factor underlying the reduced *P*_positive_ observed in OASSIS.

In addition to reducing the final positive-reactor fraction, the current branched DBD-conjugated reporter also appeared to delay signal generation. Digital RNA detection using this reporter showed delayed fluorescence development in both the conventional oil-sealed DEX droplet assay and OASSIS (Supplementary Fig. 2 and Fig. 3f). Therefore, further optimization of the reporter design will be important for improving assay performance while preserving robust signal retention within DEX droplets.

In conclusion, OASSIS establishes a new open-format digital bioassay strategy that eliminates the need for reactor sealing while preserving single-molecule sensitivity. By combining spontaneous target enrichment, signal localization, and serial sample introduction, the platform expands the design space of digital bioassays and provides a promising foundation for future amplification-free molecular diagnostics.

## Materials and Methods

### Materials and Chemicals

Polyethylene glycol (MW: 35 kDa), dextran from Leuconostoc spp. (MW: 450 - 650 kDa), TRITC-dextran (MW: 500 kDa), tetrazine(Tz)-PEG5-NHS, Benzonase® Nuclease, Dibenzocyclooctyne-N-hydroxysuccinimidyl ester (DBCO-NHS), and Amicon® Ultra-4 Centrifugal Tubes Ultracel® – 10K were purchased from Merck (Sigma-Aldrich), Germany. Fluorinert-FC40 (3M, USA), SURFLON S-386 (AGC Seimi Chemical, Japan), BSA (New England Biolabs, USA), Fomblin YLVAC25/6 (Solvay, Belgium), trans-cyclooctene (TCO)-PEG4-NHS (TCI, Japan), Ni SepharoseTM 6 Fast Flow (Cytiva, USA), Strep-Tactin® Superflow plus (Qiagen, Germany), Zeba Spin Desalting Columns (7K MWCO, 0.5 mL), HiPPR Detergent Removal Resin (Thermo Scientific, USA), Recombinant RNase Inhibitor (Takara, Japan), and cOmplete ULTRA Tablets, Mini, EDTA-free, EASYpack (Roche, Switzerland) were sourced from respective suppliers.

### Flow-channel fabrication using FRAD (femtoliter reactor array device)

FRAD were fabricated by photolithography as described previously ^32^. Flow cells comprised a FRAD and a ported top glass slide separated by double-sided adhesive tape (∼80 µm) as a spacer. The top glass was precoated with CYTOP 809M to suppress nonspecific biomolecular adsorption.

### Microscopic Imaging

Confocal and epifluorescence images were acquired using the systems described previously ^32^. Briefly, TCS SP8 (Leica) was used for confocal imaging and FRAP experiments, and IX83 (Olympus) equipped with sCMOS camera (Zyla Andor neo, Andor Technology) and LED light source (X-Cite Turbo, Excelitas Technologies) was used for epifluorescence imaging. The following filter sets were used for detection: λ_ex_ = 489 nm and λ_em_ = 508 nm for FAM. All image analyses were performed using Fiji with custom-written macros.

### Purification of tandem DBD fusion protein (DBD×2)

The expression plasmid for SUMO-DBD×2 was transformed into *Escherichia coli* Rosetta2(DE3) pLysS. Cells were cultured in LB medium at 37 °C with shaking at 200 rpm. When the OD600 reached 0.4–0.6, the culture was cooled at 4 °C for 30 min. Protein expression was induced with a final concentration of 0.5 mM IPTG, and the cells were further incubated at 21 °C with shaking at 180 rpm for 16 h. Cells were harvested by centrifugation at 5,200 × g for 10 min at 4 °C. The cell pellet was resuspended in Buffer (20 mM Tris-HCl, pH 8.0, 500 mM NaCl, 1 mM DTT, 1 mg/mL Lysozyme, 0.5 unit/µL Benzonase, and cOmplete Ultra protease inhibitor). The cells were disrupted by sonication (Amplitude 100, 1 sec on / 2 sec off, 5 min total), and the lysate was clarified by ultracentrifugation at 36,500 rpm for 60 min at 4 °C. The clarified supernatant was first applied to a Ni Sepharose 6 Fast Flow column. The column was washed with Ni-NTA Binding Buffer (20 mM Tris-HCl, pH 7.5, 500 mM NaCl, 30 mM Imidazole, 1 mM DTT), and the protein was eluted with Ni-NTA Elution Buffer (20 mM Tris-HCl, pH 7.5, 500 mM NaCl, 300 mM Imidazole, 1 mM DTT). The eluate was then applied to a StrepTactin SuperFlow plus column and washed with Wash Buffer (20 mM Tris-HCl, pH 8.0, 500 mM NaCl, 1 mM DTT). SUMO protease buffer (20 mM Tris-HCl, pH 8.0, 500 mM NaCl, 1 mM DTT, 20 nM SUMO protease (Ulp1), 0.15%(v/v) NP-40) was added to the resin, and the cleavage of the SUMO-tag was allowed to proceed overnight at 4 °C to elute the DBD×2 protein. The eluate was concentrated using an Amicon Ultra-4 (10K MWCO) centrifugal filter. Finally, NP-40 was removed using HiPPR Detergent Removal Resin following the manufacturer’s protocol.

### Preparation of the DBD-conjugated probe

Purified DBD×2 protein was buffer-exchanged to PBS–NaHCO₃ (pH 8.0; 1× PBS, pH 7.4, adjusted with 1 M NaHCO₃) using a Zeba Spin Desalting Column (7K MWCO, 0.5 mL). DBCO–NHS (dissolved in DMSO) was added to the DBD×2 solution at a 1:10 molar ratio (DBD×2:DBCO–NHS) and incubated at 4 °C for 16 h with rotation. Excess reagent was removed and the buffer was simultaneously exchanged to 1× PBS (pH 7.4; 8.1 mM Na₂HPO₄, 1.47 mM KH₂PO₄, 2.68 mM KCl, 137 mM NaCl) using a Zeba Spin Desalting Column (7K MWCO, 0.5 mL). The azide-probe dissolved in 1× TE (pH 8.0; Nippon Gene, Tokyo, Japan) was then added to the DBCO-labeled DBD×2 at a 1:3 molar ratio (DBD×2–DBCO:azide-probe), and the mixture was incubated at 4 °C for 16 h with rotation. The azide-probe used here is a branched fluorescent reporter bearing a single terminal azide. We selected the DBD×2:DBCO–NHS:azide-probe ratio based on the optimization shown in Supplementary Fig. 4 and used 1:10:3 for all assays unless otherwise noted.

### FRAP (Fluorescence Recovery After Photobleaching) experiment

For Fluorescence Recovery After Photobleaching (FRAP) analysis, several solutions were prepared. A flow cell was first infused with a blocking solution (0.5 mg/mL BSA, 0.06% (w/w) S-386 (AGC Seimi Chemical, Japan) in Buffer A (20 mM HEPES-NaOH, pH 6.8, 60 mM NaCl, 6 mM MgCl2)). The device was placed on an aluminum block on ice for 10 sec, followed by incubation at room temperature (RT) for 10 min. Subsequently, the cell was infused with a wash solution (0.06% (w/w) S-386 in Buffer A), followed by DEX Mix (0.06% (w/w) S-386, 5.5% (w/w) DEX, 0.1% (w/w) Rh-DEX in Buffer A). Next, 60 µL of PEG Mix (0.06% (w/w) S-386, 5.0% (w/w) PEG in Buffer A) was infused. Finally, the PEG Mix containing DBD-conjugated probe (0.06% (w/w) S-386, 5.0% (w/w) PEG in Buffer A) was infused into the flow cell. The device was placed on the stage of a Leica TCS SP8 confocal microscope (Leica Microsystems, Germany), and the focus was adjusted to the height of the chamber. FRAP was performed on four chambers at 25 °C using a 100x objective lens.

### Open Aqueous two-phase Separation System for Integrated Single-molecule digital bioassay (OASSIS)

The assay was performed by sequentially infusing solutions into a flow cell. First, 14 µL of blocking solution (0.5 mg/mL BSA, 0.06% (w/w) S-386 in Buffer A (20 mM HEPES-NaOH, pH 6.8, 60 mM NaCl, 6 mM MgCl2)) was infused. The device was placed on an aluminum block on ice for 10 sec, followed by incubation at room temperature (RT) for 10 min. Next, 20 µL of DEX Mix (0.06% (w/w) S-386, 5.5% (w/w) DEX, 0.01% (w/w) Rh-DEX in Buffer A) was infused. The flow cell was then flushed three times with 20 µL of PEG Mix (0.06% (w/w) S-386, 6.0% (w/w) PEG in Buffer A). After confirmation of reactor formation, 20 µL of Cas13-DBD/crRNA/Probe Mix (0.06% (w/w) S-386, 6.0% (w/w) PEG, 45 nM *Lwa*Cas13-DBD, 22.5 nM crRNA, 3 µM DBD-Probe (prepared at a 1:10:3 molar ratio of DBD:DBCO-NHS:N_3_-Probe), 2 U/µL RNase Inhibitor in Buffer A) was infused. The device was incubated at RT for 20 min. Finally, 20 µL of target RNA Mix containing 0.06% (w/w) S-386, 6.0% (w/w) PEG, 0, 0.2 fM, 1 fM, 10 fM, or 100 fM target RNA (3000 nt), and 2 U/µL RNase Inhibitor was infused into the flow cell. This infusion was repeated every 10 minutes, according to the desired number of target injections. The flow cell was observed under a microscope, and endpoint analysis was performed at 120 min.

### OASSIS for clinical sample

This study was conducted in accordance with the Declaration of Helsinki. Written informed consent was obtained from all participants for the collection of nasopharyngeal swab samples. The study protocol was approved by the Institutional Research Ethics Committee of the Faculty of Medicine, The University of Tokyo (Approval no. 2021036NI-3).

The collected nasopharyngeal swab samples were first subjected to RNase inactivation. An inactivation solution was prepared by mixing the clinical sample with denaturing agents to achieve a composition of sample, 1 M guanidine thiocyanate (GITC), and 1% (v/v) 2-mercaptoethanol (2-ME). The mixture was incubated at 95 °C for 5 min. OASSIS was performed by sequentially infusing reaction mixtures into the flow cell. First, a blocking solution containing 0.5 mg/mL BSA and 0.06% (w/w) S-386 in Buffer A (20 mM HEPES-NaOH, pH 6.8, 60 mM NaCl, 6 mM MgCl2) was infused. The flow cell was placed on an aluminum block on ice for 10 sec. Next, DEX Mix (0.06% (w/w) S-386, 5.5% (w/w) DEX, and 0.01% (w/w) Rh-DEX in Buffer A) was infused, followed by the infusion of PEG Mix (0.06% (w/w) S-386 and 6.0% (w/w) PEG in Buffer A). To initiate the reactor preparation, a mixture containing the Cas13-crRNA complex and the DBD- probe was introduced into the flow cell. This mixture consisted of 0.06% (w/w) S-386, 6.0% (w/w) PEG, 45 nM *Lwa*Cas13-DBD, 22.5 nM crRNA, 3 µM DBD-Probe (prepared at a molar ratio of 1:10:3 for DBD:DBCO-NHS:N3-Probe), and 2 U/µL RNase Inhibitor in Buffer A. The device was incubated at room temperature for 20 min. Finally, the assay was started by infusing the sample mixture containing target RNA. This final mixture contained 0.06% (w/w) S-386, 6.0% (w/w) PEG, 2 U/µL RNase Inhibitor, 0 or 300 fM Target RNA (3000 nt), and the diluted inactivated sample (resulting in final concentrations of 0.5% (v/v) nasopharyngeal swab sample, 5 mM GITC, and 0.005% (v/v) 2-ME) in Buffer A. Fluorescence microscopy images were acquired with an endpoint of 120 min.

## Author contributions

H.N. conceived the idea of the research and supervised it. Y.M. designed the experiments. Y.M., K.M., and S.N. conducted the experiments and analyzed the data. H.I., M.K., and M.N. provided technical support. H.I. and M.K. collected and prepared the specimens. H.N. and Y.M. wrote the manuscript.

## Acknowledgements

This research was supported by AMED under Grant Number JP21km0908001 (Moonshot project) and JP223fa627001 (UTOPIA project). This work was also supported by Next Generation Biomedical Measurement Research Network Program of Nakatani Foundation.

## References

(1) Basu, A. S. Digital Assays Part II: Digital Protein and Cell Assays. SLAS Technol 2017, 22 (4), 387–405. DOI: 10.1177/2472630317705681.

(2) Basu, A. S. Digital Assays Part I: Partitioning Statistics and Digital PCR. SLAS Technol 2017, 22 (4), 369–386. DOI: 10.1177/2472630317705680.

(3) Noji, H.; Minagawa, Y.; Ueno, H. Enzyme-based digital bioassay technology - key strategies and future perspectives. Lab Chip 2022, 22 (17), 3092–3109. DOI: 10.1039/d2lc00223j.

(4) Zhang, Y.; Noji, H. Digital Bioassays: Theory, Applications, and Perspectives. Anal Chem 2017, 89 (1), 92–101. DOI: 10.1021/acs.analchem.6b04290.

(5) Zhang, Y. T.; Gu, H. C.; Xu, H. Recent progress in digital immunoassay: how to achieve ultrasensitive, multiplex and clinical accessible detection? Sens Diagn 2024, 3 (1). DOI: 10.1039/d3sd00144j.

(6) Ding, X.; Mu, Y. Research and Application Progress of Digital Nucleic Acid Amplification Detection Techniques. Chinese J Anal Chem 2016, 44 (4), 512–520. DOI: 10.1016/S1872-2040(16)60918-0.

(7) Gansen, A.; Herrick, A. M.; Dimov, I. K.; Lee, L. P.; Chiu, D. T. Digital LAMP in a sample self-digitization (SD) chip. Lab Chip 2012, 12 (12), 2247–2254. DOI: 10.1039/c2lc21247a.

(8) Shen, F.; Du, W. B.; Kreutz, J. E.; Fok, A.; Ismagilov, R. F. Digital PCR on a SlipChip. Lab on a Chip 2010, 10 (20), 2666–2672. DOI: 10.1039/c004521g.

(9) Kim, S. H.; Iwai, S.; Araki, S.; Sakakihara, S.; Iino, R.; Noji, H. Large-scale femtoliter droplet array for digital counting of single biomolecules. Lab Chip 2012, 12 (23), 4986–4991. DOI: 10.1039/c2lc40632b.

(10) Rissin, D. M.; Kan, C. W.; Campbell, T. G.; Howes, S. C.; Fournier, D. R.; Song, L.; Piech, T.; Patel, P. P.; Chang, L.; Rivnak, A. J.;, et al. Single-molecule enzyme-linked immunosorbent assay detects serum proteins at subfemtomolar concentrations. Nat Biotechnol 2010, 28 (6), 595–599. DOI: 10.1038/nbt.1641.

(11) Choi, J. W.; Seo, W. H.; Kang, T.; Kang, T.; Chung, B. G. Droplet digital recombinase polymerase amplification for multiplexed detection of human coronavirus. Lab on a Chip 2023, 23 (10), 2389–2398. DOI: 10.1039/d3lc00025g.

(12) Tan, L. L.; Loganathan, N.; Agarwalla, S.; Yang, C.; Yuan, W. Y.; Zeng, J.; Wu, R. G.; Wang, W.; Duraiswamy, S. Current commercial dPCR platforms: technology and market review. Crit Rev Biotechnol 2023, 43 (3), 433–464. DOI: 10.1080/07388551.2022.2037503.

(13) Wu, C.; Garden, P. M.; Walt, D. R. Ultrasensitive Detection of Attomolar Protein Concentrations by Dropcast Single Molecule Assays. J Am Chem Soc 2020, 142 (28), 12314–12323. DOI: 10.1021/jacs.0c04331.

(14) Zhang, L.; Bai, H.; Zou, J.; Zhang, C.; Zhuang, W.; Hu, J.; Yao, Y.; Hu, W. W. Immuno-Rolling Circle Amplification (Immuno-RCA): Biosensing Strategies, Practical Applications, and Future Perspectives. Adv Healthc Mater 2024, 13 (32), e2402337. DOI: 10.1002/adhm.202402337.

(15) Akama, K.; Shirai, K.; Suzuki, S. Droplet-Free Digital Enzyme-Linked Immunosorbent Assay Based on a Tyramide Signal Amplification System. Anal Chem 2016, 88 (14), 7123–7129. DOI: 10.1021/acs.analchem.6b01148.

(16) Park, J.; Park, M.; Kim, J.; Heo, Y.; Han, B. H.; Choi, N.; Park, C.; Lee, R.; Lee, D. G.; Chung, S.;, et al. Beads- and oil-free single molecule assay with immuno-rolling circle amplification for detection of SARS-CoV-2 from saliva. Biosens Bioelectron 2023, 232, 115316. DOI: 10.1016/j.bios.2023.115316.

(17) Xie, Y.; Li, H.; Tang, Y.; Lian, X.; Dai, L.; Tian, S. Droplet-free and enzyme-free digital immunoassay based on fluorescent microspheres for protein detection. Sensors and Actuators B: Chemical 2023, 396. DOI: 10.1016/j.snb.2023.134547.

(18) Wang, S. Q.; Lifson, M. A.; Inci, F.; Liang, L. G.; Sheng, Y. F.; Demirci, U. Advances in addressing technical challenges of point-of-care diagnostics in resource-limited settings. Expert Rev Mol Diagn 2016, 16 (4), 449–459. DOI: 10.1586/14737159.2016.1142877.

(19) Park, J.; Han, D. H.; Park, J. K. Towards practical sample preparation in point-of-care testing: user-friendly microfluidic devices. Lab on a Chip 2020, 20 (7), 1191–1203. DOI: 10.1039/d0lc00047g.

(20) Liu, D.; Wang, Y.; Li, X.; Li, M.; Wu, Q.; Song, Y.; Zhu, Z.; Yang, C. Integrated microfluidic devices for in vitro diagnostics at point of care. Aggregate 2022, 3 (5). DOI: 10.1002/agt2.184.

(21) Shinoda, H.; Iida, T.; Makino, A.; Yoshimura, M.; Ishikawa, J.; Ando, J.; Murai, K.; Sugiyama, K.; Muramoto, Y.; Nakano, M.;, et al. Automated amplification-free digital RNA detection platform for rapid and sensitive SARS-CoV-2 diagnosis. Commun Biol 2022, 5 (1), 473. DOI: 10.1038/s42003-022-03433-6.

(22) Akama, K.; Iwanaga, N.; Yamawaki, K.; Okuda, M.; Jain, K.; Ueno, H.; Soga, N.; Minagawa, Y.; Noji, H. Wash- and Amplification-Free Digital Immunoassay Based on Single-Particle MotionAnalysis. ACS Nano 2019, 13 (11), 13116–13126. DOI: 10.1021/acsnano.9b05917.

(23) Witters, D.; Knez, K.; Ceyssens, F.; Puers, R.; Lammertyn, J. Digital microfluidics-enabled single-molecule detection by printing and sealing single magnetic beads in femtoliter droplets. Lab Chip 2013, 13 (11), 2047–2054. DOI: 10.1039/c3lc50119a.

(24) Nakatani, N.; Sakuta, H.; Hayashi, M.; Tanaka, S.; Takiguchi, K.; Tsumoto, K.; Yoshikawa, K. Specific Spatial Localization of Actin and DNA in a Water/Water Microdroplet: Self-Emergence of a Cell-Like Structure. Chembiochem 2018, 19 (13), 1370–1374. DOI: 10.1002/cbic.201800066.

(25) Strulson, C. A.; Molden, R. C.; Keating, C. D.; Bevilacqua, P. C. RNA catalysis through compartmentalization. Nat Chem 2012, 4 (11), 941–946. DOI: 10.1038/nchem.1466.

(26) Mizuuchi, R.; Ichihashi, N. Translation-coupled RNA replication and parasitic replicators in membrane-free compartments. Chem Commun (Camb) 2020, 56 (87), 13453–13456. DOI: 10.1039/d0cc06606k.

(27) Diamond, A. D.; Hsu, J. T. Protein partitioning in PEG/dextran aqueous two - phase systems. AIChE Journal 2004, 36 (7), 1017–1024. DOI: 10.1002/aic.690360707.

(28) Zhang, Y.; Kojima, T.; Kim, G. A.; McNerney, M. P.; Takayama, S.; Styczynski, M. P. Protocell arrays for simultaneous detection of diverse analytes. Nat Commun 2021, 12 (1), 5724. DOI: 10.1038/s41467-021-25989-3.

(29) Ahmed, T.; Verma, A.; Patterson, A. T.; Styczynski, M. P.; Takayama, S. Compositional Dependence of DNA Partitioning in a Poly(Ethylene Glycol)‒Ficoll Aqueous Two-Phase System. Chemistry 2024, 6 (6), 1680–1691. DOI: 10.3390/chemistry6060102.

(30) Zha, Q.; Luo, Y.; Liu, C.; Xu, T. Integrated phase separation in microliter droplets for ultratrace-enriching biomarker analysis. Lab Chip 2024, 24 (6), 1775–1781. DOI: 10.1039/d3lc00953j.

(31) Lv, H.; Duan, X.; Han, Z.; Yu, H.; Liu, B. Quencher-free fluorescent assays by controlled DNA partitioning in the aqueous two-phase system with crowding-enhanced kinetics. Biosens Bioelectron 2024, 246, 115864. DOI: 10.1016/j.bios.2023.115864.

(32) Minagawa, Y.; Nakata, S.; Date, M.; Ii, Y.; Noji, H. On-Chip Enrichment System for Digital Bioassay Based on Aqueous Two-Phase System. ACS Nano 2023, 17 (1), 212–220. DOI: 10.1021/acsnano.2c06007.

(33) Suwannarangsee, S.; Moulis, C.; Potocki-Veronese, G.; Monsan, P.; Remaud-Simeon, M.; Chulalaksananukul, W. Search for a dextransucrase minimal motif involved in dextran binding. FEBS Lett 2007, 581 (24), 4675–4680. DOI: 10.1016/j.febslet.2007.08.062.

(34) Choi, J. W.; Jung, D.; Park, Y. M.; Bae, N. H.; Lee, S. J.; Rho, D.; Chung, B. G.; Lee, K. G. Microinjection molded microwell array-based portable digital PCR system for the detection of infectious respiratory viruses. Nano Converg 2025, 12 (1), 16. DOI: 10.1186/s40580-025-00482-5.

(35) Honda, S.; Minagawa, Y.; Noji, H.; Tabata, K. V. Multidimensional Digital Bioassay Platform Based on an Air-Sealed Femtoliter Reactor Array Device. Anal Chem 2021, 93 (13), 5494–5502. DOI: 10.1021/acs.analchem.0c05360.

(36) Yaginuma, H.; Ohtake, K.; Akamatsu, T.; Noji, H.; Tabata, K. A microreactor sealing method using adhesive tape for digital bioassays. Lab on a Chip 2022, 22 (10), 2001–2010. DOI: 10.1039/d2lc00065b.

(37) Tian, T.; Shu, B.; Jiang, Y.; Ye, M.; Liu, L.; Guo, Z.; Han, Z.; Wang, Z.; Zhou, X. An Ultralocalized Cas13a Assay Enables Universal and Nucleic Acid Amplification-Free Single-Molecule RNA Diagnostics. ACS Nano 2021, 15 (1), 1167–1178. DOI: 10.1021/acsnano.0c08165.

(38) Shinoda, H.; Taguchi, Y.; Nakagawa, R.; Makino, A.; Okazaki, S.; Nakano, M.; Muramoto, Y.; Takahashi, C.; Takahashi, I.; Ando, J.;, et al. Amplification-free RNA detection with CRISPR-Cas13. Commun Biol 2021, 4 (1), 476. DOI: 10.1038/s42003-021-02001-8.

(39) Kellner, M. J.; Koob, J. G.; Gootenberg, J. S.; Abudayyeh, O. O.; Zhang, F. SHERLOCK: nucleic acid detection with CRISPR nucleases. Nat Protoc 2019, 14 (10), 2986–3012. DOI: 10.1038/s41596-019-0210-2.

(40) East-Seletsky, A.; O’Connell, M. R.; Burstein, D.; Knott, G. J.; Doudna, J. A. RNA Targeting by Functionally Orthogonal Type VI-A CRISPR-Cas Enzymes. Mol Cell 2017, 66 (3), 373–383 e373. DOI: 10.1016/j.molcel.2017.04.008.

(41) Minagawa, Y.; Yabuta, M.; Su’etsugu, M.; Noji, H. Self-growing protocell models in aqueous two-phase system induced by internal DNA replication reaction. Nat Commun 2025, 16 (1), 1522. DOI: 10.1038/s41467-025-56172-7.

(42) Kalnina, L.; Mateu-Regue, A.; Oerum, S.; Hald, A.; Gerstoft, J.; Oerum, H.; Nielsen, F. C.; Iversen, A. K. N. A simple, safe and sensitive method for SARS-CoV-2 inactivation and RNA extraction for RT-qPCR. APMIS 2021, 129 (7), 393–400. DOI: 10.1111/apm.13123.

