## SupplementalFigures for "Open Digital Bioassays Enabled by Aqueous Two-Phase Microreactor Arrays"

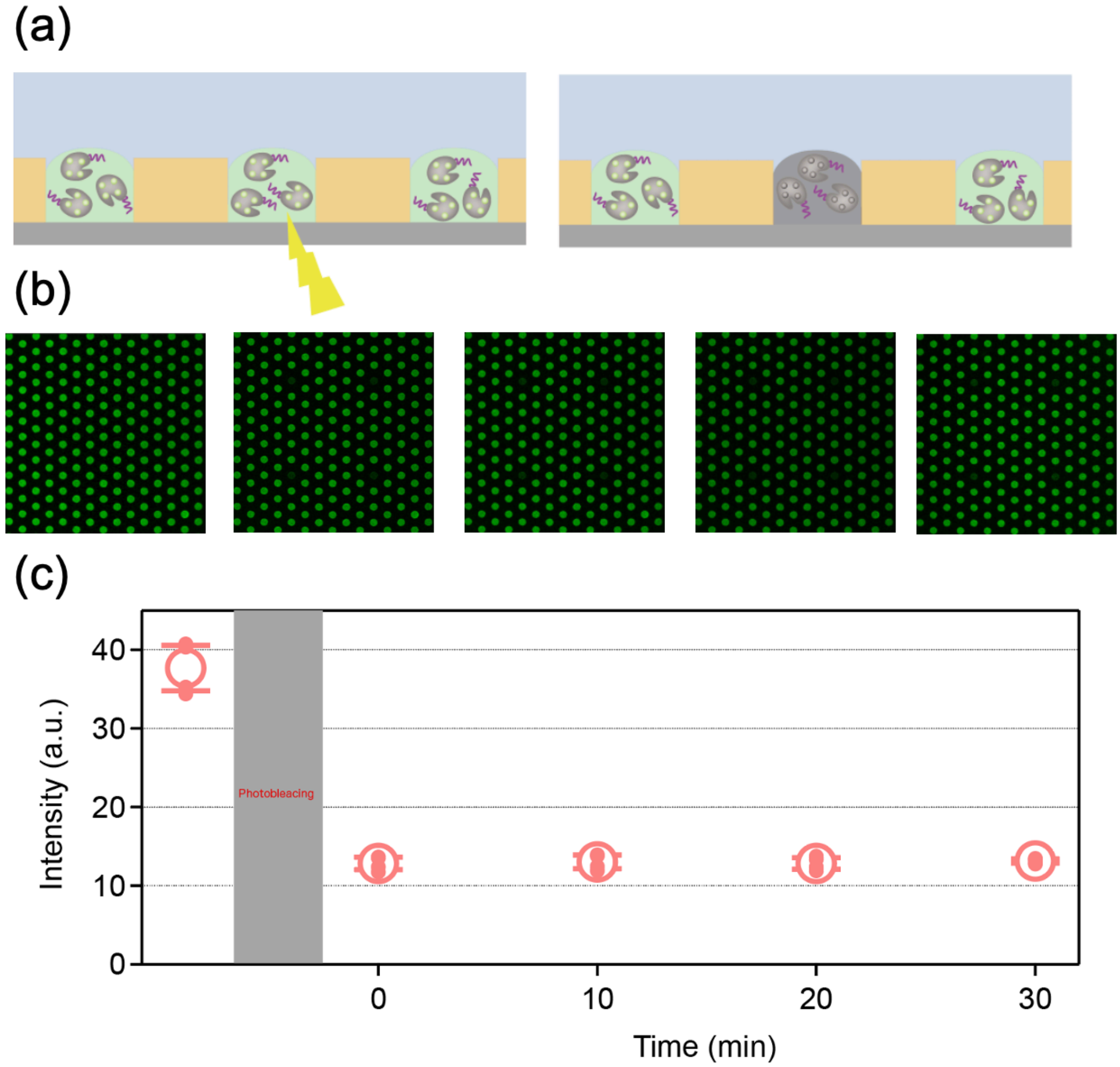


Supplementary Figure 1. FRAP analysis of Alexa Fluor 488–labeled Cas13-DBD fusion protein.

(a) Schematic of the fluorescence recovery after photobleaching (FRAP) experiment targeting Alexa Fluor 488–labeled Cas13-DBD fusion protein enriched in DEX droplets. (b) Time-lapse fluorescence images of the reactor array before and after photobleaching. (c) Quantitative analysis of fluorescence recovery. Similar to the DBD-conjugated probe (main Fig. 2), the Cas13-DBD protein shows no fluorescence recovery after photobleaching, confirming its robust confinement within the DEX droplets without detectable diffusion between reactors.

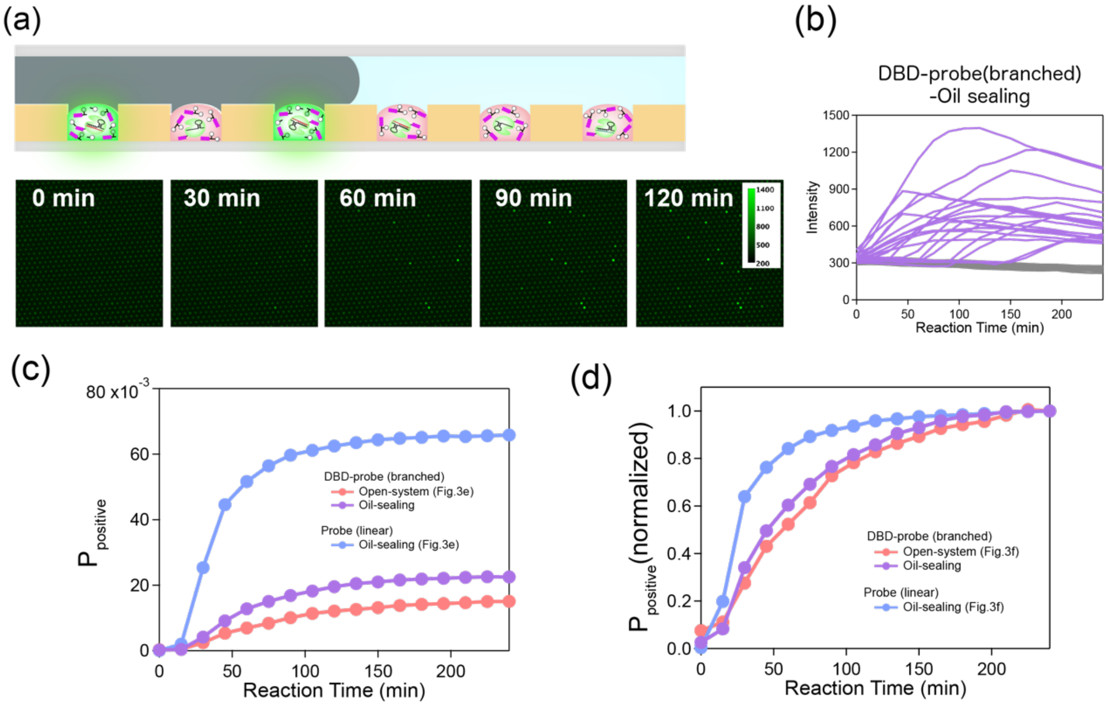


Supplementary Figure 2. Digital RNA detection with the DBD-conjugated branched reporter in the oil-sealed DEX droplet system.

(a) Schematic and time-lapse fluorescence images of Cas13 digital RNA detection performed in the oil-sealed DEX droplet system. (b) Representative fluorescence time-courses of individual reactors classified as positive (purple) or negative (gray). (c) Temporal evolution of the probability of positive reactors ($P_{\text{positive}}$) for the oil-sealed system using the conventional linear reporter (blue; main Fig. 3e), the oil-sealed system using the DBD-conjugated branched reporter (purple), and the open system using the DBD-conjugated branched reporter (red; main Fig. 3e). (d) Normalized $P_{\text{positive}}$ time-courses for the same assay configurations as in (c). The oil-sealed assay using the DBD-conjugated branched reporter showed a lower final $P_{\text{positive}}$ than the oil-sealed assay using the conventional linear reporter, while its normalized time-course was similar to that observed in the open system. These results suggest that the reduced final positive-reactor fraction observed in the open system is mainly associated with the branched reporter design rather than the open-format configuration itself.

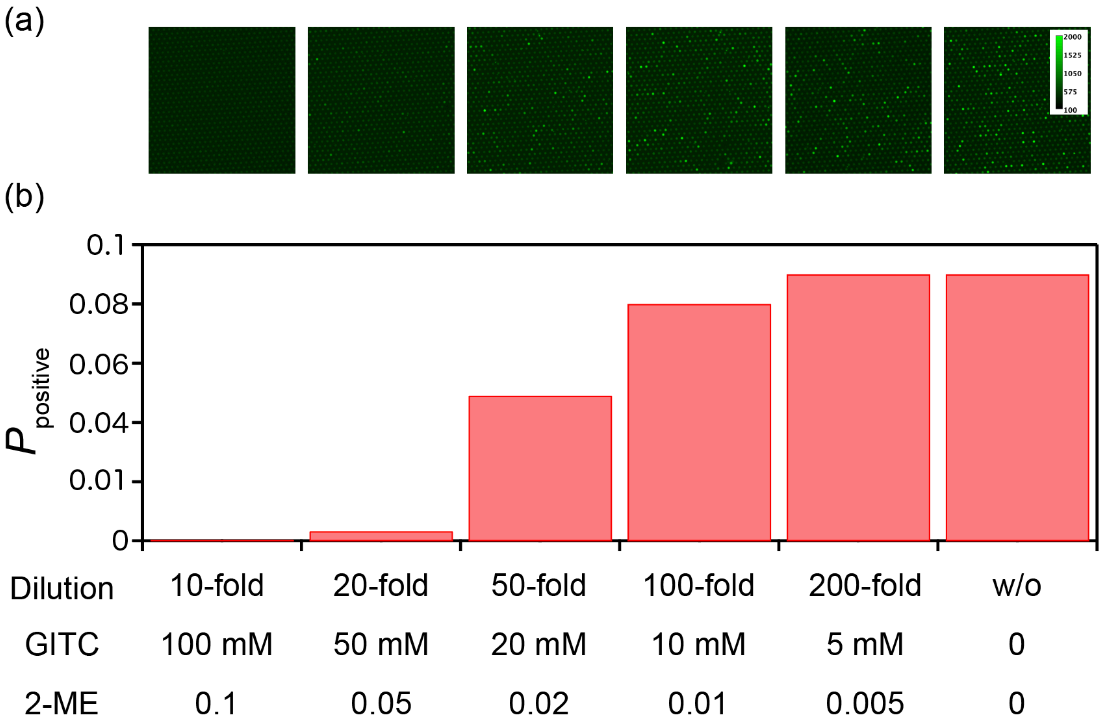


Supplementary Figure 3. Cas13a activity in the presence of chemical denaturants.

The effect of RNase inactivation reagents was investigated: guanidinium thiocyanate (GITC) and 2-mercaptoethanol (2-ME). The probability of positive reactors ($P_{positive}$) was measured under various dilution factors of the inactivation mixture. While high concentrations of GITC and 2-ME inhibit the enzymatic activity, a 200-fold dilution (resulting in 5 mM GITC and 0.005% 2-ME) restores the signal to levels comparable to the control (w/o denaturants).

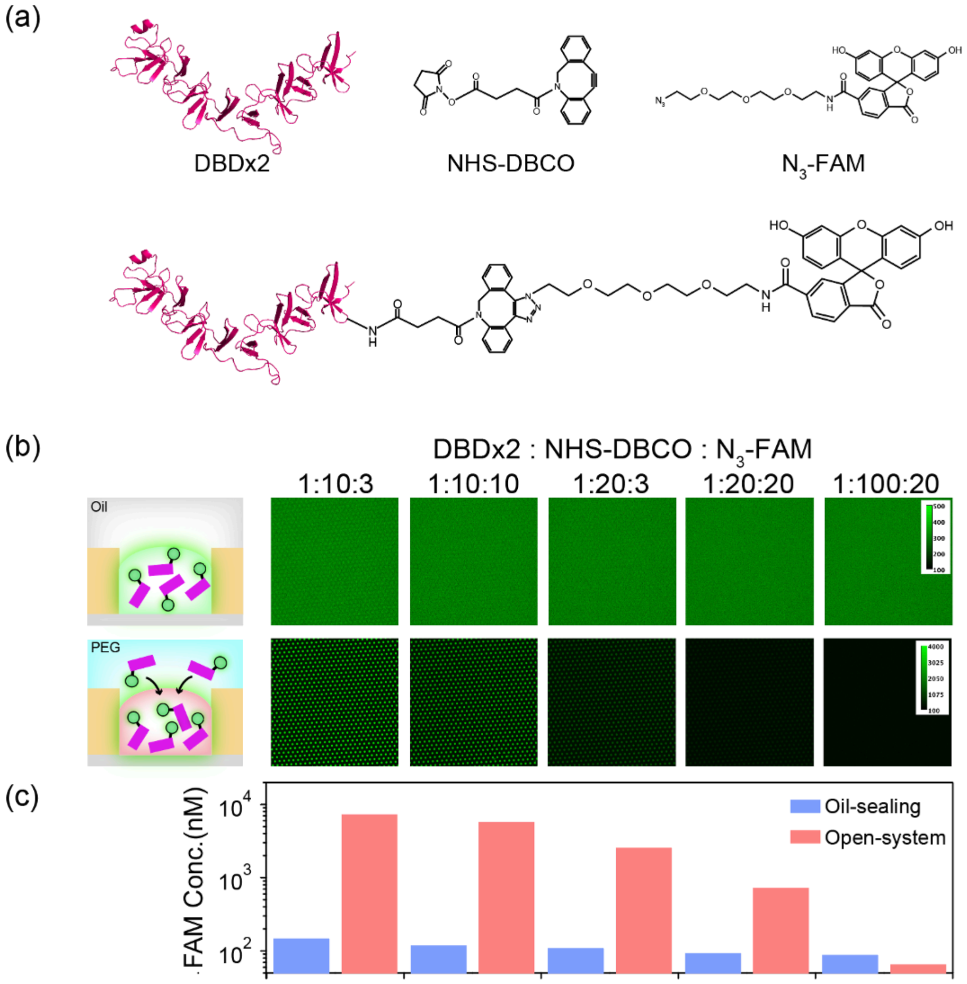


Supplementary Figure 4. Optimization of the conjugation ratio for the DBD-conjugated branched probe.

(a) Reaction scheme for the synthesis of the DBD-conjugated probe. The tandem dextran-binding domain (DBD×2) was first labeled with a heterobifunctional linker (NHS-DBCO), followed by conjugation with the azide-functionalized branched fluorescent reporter (N3-FAM) via copper-free click chemistry. (b) Fluorescence images of DEX reactors loaded with DBD-probes synthesized at various molar ratios of reagents (DBD×2 : NHS-DBCO : N3-FAM). Images compare the reactor arrays in the conventional oil-sealed system (top row) and the open ATPS system (bottom row). (c) Quantification of the estimated concentration of the conjugated probe within the DEX reactors. The red bars represent the open-system (OASSIS), where the probe is enriched via the DBD moiety, while blue bars represent the oil-sealed system. The molar ratio of 1:10:3 yielded the highest effective concentration in the open-system reactors, demonstrating maximal enrichment efficiency. Based on this result, the 1:10:3 ratio was adopted for all experiments in this study.

Supplementary Table 1 RNA, crRNA, and self-quenched probe sequences used in this study

| | |
|---|---|
| Target RNA | GGGUAACAUCACUAGGUUUCAAACUUUACUUGCUUUACAUAGAAGUUA<br>UUUGACUCCUGGUGAUUCUUCUUCAGGUUGGACAGCUGGUGCUGCAGC<br>UUAUUAUGUGGGUUAUCUUCAACCUAGGGAGAACAGCAAGAAGCACGA<br>GAAGUACAAGAUCCGCGAGUACUAUCACAAGAUCAUCGGCCGGAAGAA<br>CGACAAAGAGAACUUCGCCAAGAUUAUCUACGAAGAGAUCCAGAACGU<br>GAACAACAUCAAAGAGCUGAUUGAGAAGAUCCCCGACAUGUCUGAGCU<br>GAAGAAAAGCCAGGUGUUCUACAAGUACUACCUGGACAAAGAGGAACU<br>GAACGACAAGAAUAUUAAGUACGCCUUCUGCCACUUCGUGGAAAUCGA<br>GAUGUCCCAGCUGCUGAAAAACUACGUGUACAAGCGGCUGAGCAACAU<br>CAGCAACGAUAAGAUCAAGCGGAUCUUCGAGUACCAGAAUCUGAAAAA<br>GCUGAUCGAAAACAAACUGCUGAACAAGCUGGACACCUACGUGCGGAA<br>CUGCGGCAAGUACAACUACUAUCUGCAAGUGGGCGAGAUCGCCACCUCC<br>GACUUUAUCGCCCGGAACCGGCAGAACGAGGCCUUCCUGAGAAACAUCA<br>UCGGCGUGUCCAGCGUGGCCUACUUCAGCCUGAGGAACAUCCUGGAAAC<br>CGAGAACGAGAACGAUAUCACCGGCCGGAUGCGGGGCAAGACCGUGAA<br>GAACAACAAGGGCGAAGAGAAAUACGUGUCCGGCGAGGUGGACAAGAU<br>CUACAAUGAGAACAAGCAGAACGAAGUGAAAGAAAAUCUGAAGAUGUU<br>CUACAGCUACGACUUCAACAUGGACAACAAGAACGAGAUCGAGGACUU<br>CUUCGCCAACAUCGACGAGGCCAUCAGCAGCAUCAGACACGGCAUCGUG<br>CACUUCAACCUGGAACUGGAAGGCAAGGACAUCUUCGCCUUCAAGAAU<br>AUCGCCCCCAGCGAGAUCUCCAAGAAGAUGUUUCAGAACGAAAUCAACG<br>AAAAGAAGCUGAAGCUGAAAAUCUUCAAGCAGCUGAACAGCGCCAACG<br>UGUUCAACUACUACGAGAAGGAUGUGAUCAUCAAGUACCUGAAGAAUA<br>CCAAGUUCAACUUCGUGAACAAAAACAUCCCCUUCGUGCCCAGCUUCAC<br>CAAGCUGUACAACAAGAUUGAGGACCUGCGGAAUACCCUGAAGUUUUU<br>UUGGAGCGUGCCCAAGGACAAAGAAGAGAAGGACGCCCAGAUCUACCU<br>GCUGAAGAAUAUCUACUACGGCGAGUUCCUGAACAAGUUCGUGAAAAA<br>CUCCAAGGUGUUCUUUAAGAUCACCAAUGAAGUGAUCAAGAUUAACAA<br>GCAGCGGAACCAGAAAACCGGCCACUACAAGUAUCAGAAGUUCGAGAA<br>CAUCGAGAAAACCGUGCCCGUGGAAUACCUGGCCAUCAUCCAGAGCAGA<br>GAGAUGAUCAACAACCAGGACAAAGAGGAAAAGAAUACCUACAUCGAC<br>UUUAUUCAGCAGAUUUUCCUGAAGGGCUUCAUCGACUACCUGAACAAG<br>AACAAUCUGAAGUAUAUCGAGAGCAACAACAACAAUGACAACAACGAC<br>AUCUUCUCCAAGAUCAAGAUCAAAAAGGAUAACAAAGAGAAGUACGAC<br>AAGAUCCUGAAGAACUAUGAGAAGCACAAUCGGAACAAAGAAAUCCCU<br>CACGAGAUCAAUGAGUUCGUGCGCGAGAUCAAGCUGGGGAAGAUUCUG<br>AAGUACACCGAGAAUCUGAACAUGUUUUACCUGAUCCUGAAGCUGCUG<br>AACCACAAAGAGCUGACCAACCUGAAGGGCAGCCUGGAAAAGUACCAG<br>UCCGCCAACAAAGAAGAAACCUUCAGCGACGAGCUGGAACUGAUCAACC<br>UGCUGAACCUGGACAACAACAGAGUGACCGAGGACUUCGAGCUGGAAG<br>CCAACGAGAUCGGCAAGUUCCUGGACUUCAACGAAAACAAAAUCAAGG<br>ACCGGAAAGAGCUGAAAAAGUUCGACACCAACAAGAUCUAUUUCGACG<br>GCGAGAACAUCAUCAAGCACCGGGCCUUCUACAAUAUCAAGAAAUACG<br>GCAUGCUGAAUCUGCUGGAAAAGAUCGCCGAUAAGGCCAAGUAUAAGA |

| | |
|---|---|
| | UCAGCCUGAAAGAACUGAAAGAGUACAGCAACAAGAAGAAUGAGAUUG<br>AAAAGAACUACACCAUGCAGCAGAACCUGCACCGGAAGUACGCCAGACC<br>CAAGAAGGACGAAAAGUUCAACGACGAGGACUACAAAGAGUAUGAGAA<br>GGCCAUCGGCAACAUCCAGAAGUACACCCACCUGAAGAACAAGGUGGAA<br>UUCAAUGAGCUGAACCUGCUGCAGGGCCUGCUGCUGAAGAUCCUGCACC<br>GGCUCGUGGGCUACACCAGCAUCUGGGAGCGGGACCUGAGAUUCCGGCU<br>GAAGGGCGAGUUUCCCGAGAACCACUACAUCGAGGAAAUUUUCAAUUU<br>CGACAACUCCAAGAAUGUGAAGUACAAAAGCGGCCAGAUCGUGGAAAA<br>GUAUAUCAACUUCUACAAAGAACUGUACAAGGACAAUGUGGAAAAGCG<br>GAGCAUCUACUCCGACAAGAAAGUGAAGAAACUGAAGCAGGAAAAAAA<br>GGACCUGUACAUCCGGAACUACAUUGCCCACUUCAACUACAUCCCCCAC<br>GCCGAGAUUAGCCUGCUGGAAGUGCUGGAAAACCUGCGGAAGCUGCUG<br>UCCUACGACCGGAAGCUGAAGAACGCCAUCAUGAAGUCCAUCGUGGACA<br>UUCUGAAAGAAUACGGCUUCGUGGCCACCUUCAAGAUCGGCGCUGACA<br>AGAAGAUCGAAAUCCAGACCCUGGAAUCAGAGAAGAUCGUGCACCUGA<br>AGAAUCUGAAGAAAAAGAAACUGAUGACCGACCGGAACAGCGAGGAAC<br>UGUGCGAACUCGUGAAAGUCAUGUUCGAGUACAAGGCCCUGGAAUAAG<br>CGGCCGCACUCGAGGCCCGAAAGGAAGCUGAGUUGGCUGCUGCCACCGC<br>UGAGCAAUAA |
| crRNA | GAUUUAGACUACCCCAAAAACGAAGGGGACUAAAACGCAGCACCAGCU<br>GUCCAACCUGAAGAAG |
| FAM-AU-BHQ1 | 6FAM-taAUgc-BHQ1* |
| Branched reporter | 5′BHQ1 - taAUgc(FLdT)tctgacgtagaaacgaggc - 3′Azide |

* Uppercase and lowercase letters represent RNA and DNA, respectively
